# Sampling the Training Trajectory of Vision Models Reveals a Mismatch with Human Visual Development

**DOI:** 10.64898/2026.11.24.768810

**Authors:** Michela Proietti, Domenic Bersch, Timothy Schaumlöffel, Diego García Cerdas, Antonia Franaszek-Traczewska, Hannah Elisabeth Zwad, Siying Xie, Marlena Baldauf, Teresa Sylvester, Christina Maria Schätz, Stefanie Höhl, Moritz Köster, Radoslaw Martin Cichy, Gemma Roig

**Affiliations:** Goethe University Frankfurt The Hessian Center for AI; Freie Universität Berlin; University of Regensburg; University of Vienna

## Abstract

We compare time-resolved EEG representational geometry across seven age groups, from 6 months to adulthood, with representations sampled throughout training from supervised and self-supervised ResNet-50 and ViT-B/16. Previous model-brain comparisons mostly used adult brain recordings and untrained or fully trained networks. We find that successive training stages do not systematically progress from child-to adult-like neural representations: alignment with every age group from 4 years onward emerges early in training and remains stable. Training adds substantial alignment beyond the untrained architecture, similarly at every age. Finally, categorical structure and spatial frequency each explain a stable share of the neural geometry at every age, with apparent age-related increases in categorical organization reflecting increased EEG reliability. Because standard training on full-resolution images aligns with all ages early, and spatial frequency, which depends on maturing visual acuity, strongly shapes the neural geometry, developmentally structured visual input may help models follow human visual development.

## 1 Introduction

Deep neural networks are widely used computational models of visual processing, with representations from trained networks predicting neural responses along the primate and human ventral visual stream [10; 23; 20; 24]. However, different models may produce similar alignment scores with adult neural data despite relying on substantially different internal representations [5]. Endpoint-based model-brain comparisons ask which fully trained model best matches the mature visual system, potentially overlooking developmental changes in brain representations. Human visual representations change substantially from infancy through childhood and adolescence, both in their spatial organiza-tion and temporal dynamics [6; 1; 22; 16; 17], despite category-selective structure emerging early in life [6; 17]. Artificial networks also undergo substantial representational change during optimization: intermediate training states first align with early sensory responses and later with higher-level and prefrontal responses [18]. However, this progression was characterized against the mature cortical and temporal hierarchy, rather than across human developmental stages. Thus, whether successive stages of model training correspond to successive stages of human visual development remains unknown. Here we address this gap using time-resolved EEG from seven age groups, spanning 6 months to adulthood, during viewing of still images. We compare neural data with ResNet-50 [9] and ViT-B/16 [7] models trained with supervised and DINO [3] self-supervised objectives. Rather than comparing untrained and fully trained networks [17], we sample representations across training progression and network depth, asking whether successive stages of model training preferentially correspond to successive stages of human visual development.

## 2 Methods

### Dataset and preprocessing

We analyzed EEG recordings from 237 participants across seven age groups: 6-month-olds (*N* = 37), 12-month-olds (*N* = 34), 4-year-olds (*N* = 36), 6-year-olds (*N* = 37), 8-year-olds (*N* = 35), 12-year-olds (*N* = 38), and adults (*N* = 20; mean age 24.8 ± 3.95 years). The dataset has been previously used to characterize developmental changes in visual representations [12] and model-brain correspondence in older children and adults [2]. Participants repeatedly viewed 48 colored images from four categories (faces, objects, scenes, and body parts), divided into four 12-image subsets. Each participant viewed one or more subsets. Participants were retained for a subset only when at least 3 repetitions per image were available. EEG epochs were baseline-corrected separately for each trial and channel and downsampled to 100 Hz.

### Models

We used ResNet-50 [9] and ViT-B/16 [7], trained on ImageNet-1K [19] with supervised classification or DINO [3] self-supervision, using three random seeds per configuration. We sampled the untrained network and 25 uniformly spaced model states. To relate training to development, we mapped training progress linearly onto a nominal age from 0-24 years and additionally saved the checkpoints whose nominal age matched each age group. Features were extracted at six depths: the convolutional stem, four residual stages, and final pooling layer for ResNet-50; five encoder blocks and the final normalization layer for ViT-B/16, using the CLS token.

### Representational dissimilarity matrices

EEG RDMs were computed per participant, stimulus subset, and time point with the Crossnobis distance [21; 8]. The trials of each image were split into up to five folds, each fold was noise-normalized with a Ledoit-Wolf covariance [11] estimated from the remaining folds, and distances were computed between independent folds (averaged over fold pairs and 20 random splits). Unlike correlation distance, crossnobis is not biased by noise [21], which matters when comparing age groups with very different signal-to-noise ratios. Model RDMs were computed as 1 − *r*, with *r* the Pearson correlation between feature vectors.

### Alignment and noise ceilings

Model-brain correspondence was quantified by Spearman correlation between each participant’s EEG RDM with the model RDM, averaged across subsets. Neural reliability was estimated for each age group using the lower noise ceiling, i.e. the correlation between each participant’s RDM and the mean RDM of the other participants [15]. Time windows with reliable EEG geometry were defined per age group as post-stimulus clusters where the lower ceiling exceeded zero (one-sided sign-flip cluster test, *p* < 0.05). The 6- and 12-month-olds showed no such cluster and were excluded from comparisons across ages. To compare ages, alignment averaged over a common reliable window (*≈* 0.14 −1.0 s) was divided by the mean lower ceiling over the same window (normalized alignment). Uncertainty was estimated with 10000 bootstrap resamples of participants within each age group. To isolate the alignment carried by learned features, the final-checkpoint RDM was residualized against the untrained RDM of the same layer before correlation with the EEG (semi-partial Spearman correlation), and normalized in the same way.

### Developmental correspondence of training

For each model, we averaged alignment within the common reliable window across all age groups and built a matrix *M* of *n* age groups ×*n* age-matched checkpoints. After dividing each row by the age’s lower ceiling and subtracting each row and column’s mean 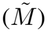, we computed the diagonal score:

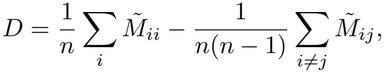

which is positive when each age group aligns best with its matched checkpoint. Significance was assessed against all *n*! permutations of the checkpoint–age pairing, and uncertainty with 1000 participant bootstrap resamples. Depth was chosen per model by the best mean rank of alignment across ages over trained checkpoints, which is invariant to checkpoint-age-pairing permutations.

### Representational content

We computed Spearman correlation between each participant’s EEG RDM and (1) a categorical RDM (0 for within- and 1 for between-category pairs), and (2) four low-level RDMs (pixel intensity, color histogram, Gabor energy, and spatial frequency). We averaged alignment within the common reliable window and normalized it by the lower ceiling. The categorical and low-level predictors were nearly orthogonal (all |*r*| < 0.09).

## 3 Results

### Fully trained models align with EEG geometry from 4 years onward, to a similar degree relative to data reliability

We first asked how alignment with fully-trained models varies across human development, neural processing time, and layer depth. In all age groups from 4 years onward, alignment emerged around 100 ms after stimulus onset and persisted throughout the epoch, with deeper representations showing stronger sustained correspondence (Figure 1; other models in Appendix Figure 2). This is consistent with prior evidence that visual DNNs capture rapidly emerging neural representations [4]. The EEG geometry of 6- and 12-month-olds was not reliable in this paradigm, and no model aligned with it significantly (Appendix Figure 2). Raw alignment was higher in older age groups, but so was the noise ceiling. Normalized by the ceiling, fully-trained models capture 40 − 80% of the explainable geometry at every age, and no pair of ages differed significantly for any model (FDR *q* < .05; Appendix Figure 3). The age differences in raw alignment therefore reflect the reliability of the EEG rather than a closer match with mature representations.

**Figure 1:**
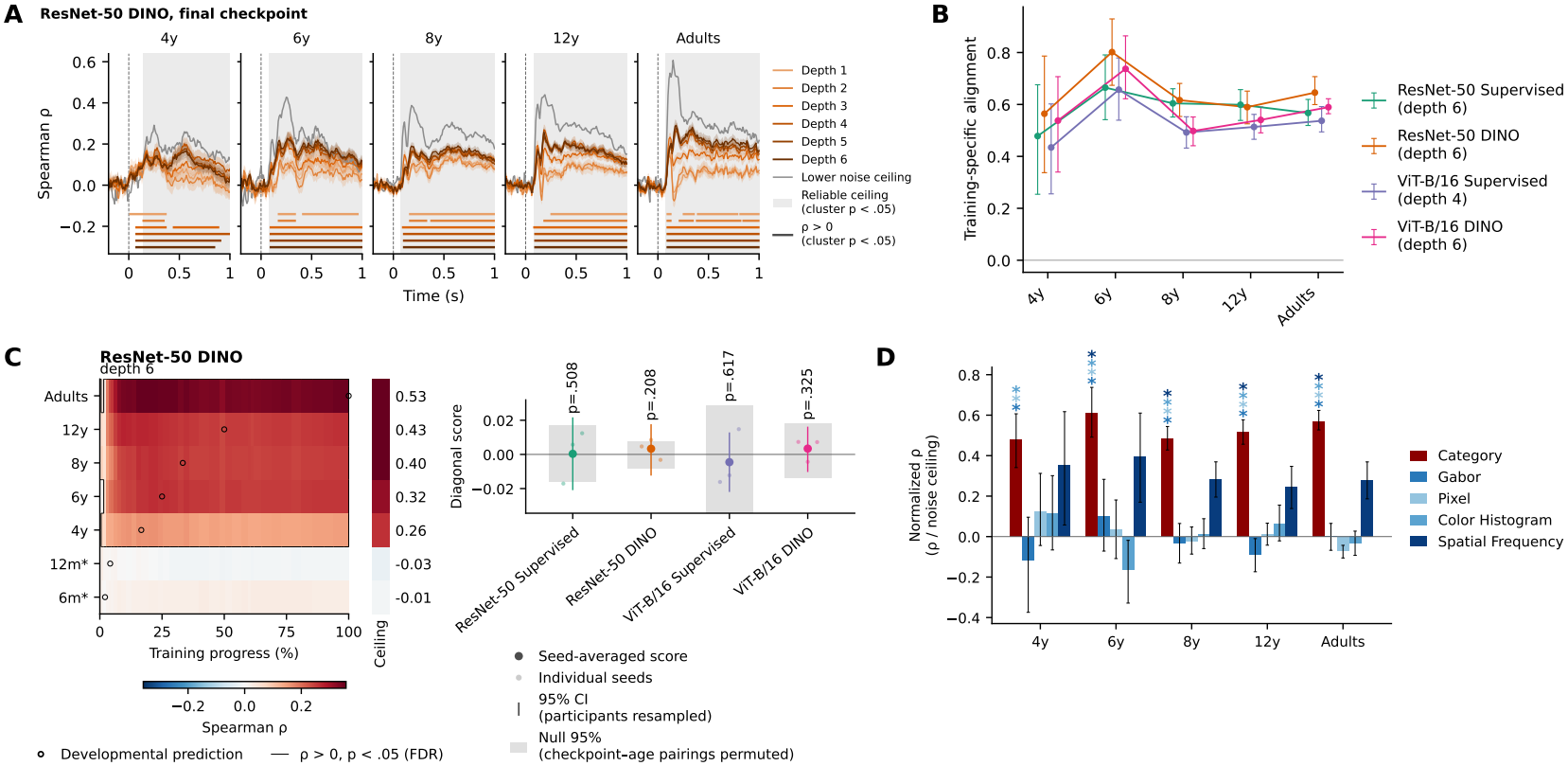
**(A)** Time-resolved RSA between EEG and model RDMs for fully trained ResNet-50 DINO (mean ±SEM over participants, one line per depth). Grey line: lower noise ceiling. Bars: *ρ* > 0 (sign-flip cluster test, *p* < .05). **(B)** Alignment uniquely attributable to learned features, normalized by the lower ceiling (common reliable window; bootstrap 95% CI). All values exceed zero (sign-flip test, FDR *q* < .05); no model shows a trend across ages (Holm *p* = 1). **(C)** Left: alignment of ResNet-50 DINO with each age group across training (common reliable window); circles mark the checkpoint whose nominal age matches each age group, while asterisks mark ages without a reliable noise ceiling. Right: diagonal score per model (*p* uncorrected, none significant after Holm correction across models). **(D)** Correlation of the EEG geometry with categorical and low-level predictors, normalized by the noise ceiling (common reliable window; bootstrap 95% CI). Stars: category > feature (one-sided sign-flip test, FDR *q* < .05).

**Figure 2:**
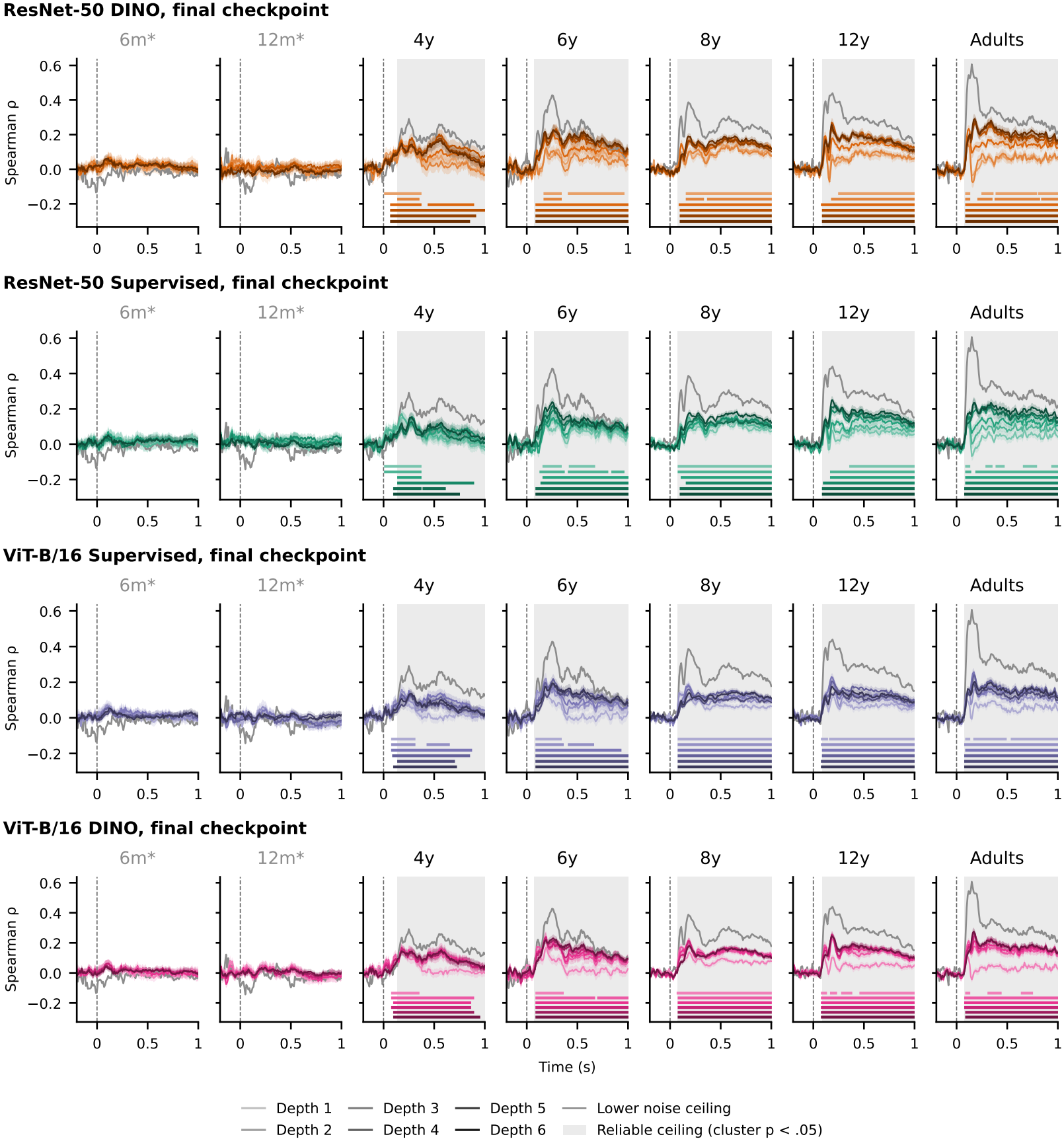
Time-resolved RSA between EEG and fully-trained model RDMs (mean ± SEM over participants, one line per depth). Grey line: lower noise ceiling. Bars: *ρ* > 0 (sign-flip cluster test, *p* < .05).

**Figure 3:**
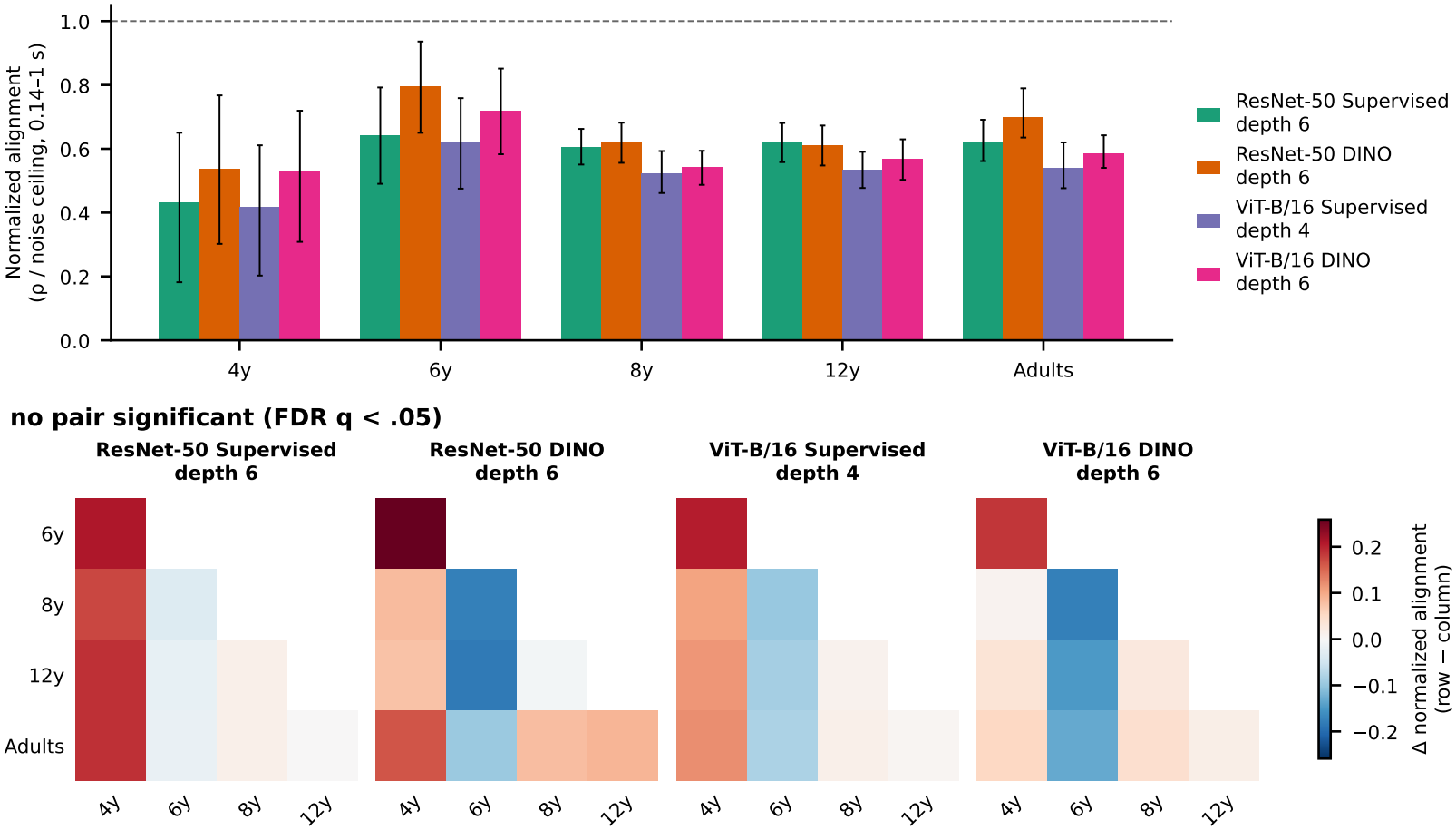
Top: normalized alignment (mean *ρ* / mean lower noise ceiling, 0.14 − 1.0 s) of each fully trained model with each age group with a reliable ceiling, at the depth used in Figure 1C. Error bars: bootstrap 95% CI over participants (10,000 resamples within each age group). Dashed line: noise ceiling. Bottom: pairwise differences between age groups (row minus column); no pair is significant (two-sided bootstrap test, FDR *q* < .05 within each model).

**Figure 4:**
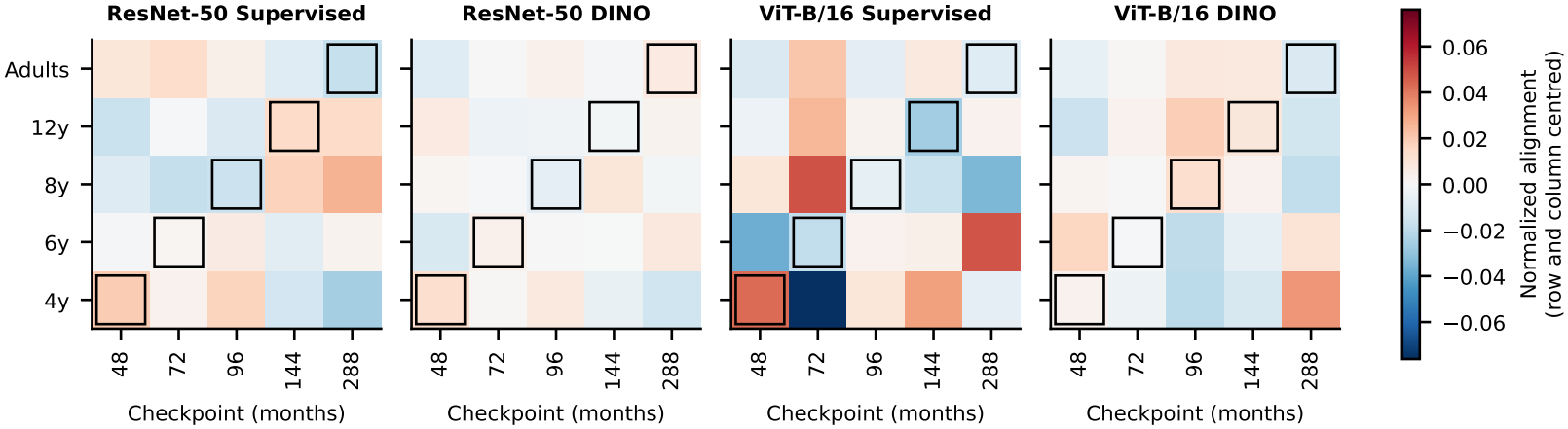
Normalized alignment of each age group with each age-matched checkpoint (rows divided by the age group’s lower noise ceiling, then row and column means removed), at the depth used in Figure 1C. Outlined cells: matched pairs (developmental prediction). The diagonal score is the mean of the outlined cells minus the mean of the others.

### Learning adds substantial alignment at every age beyond architecture alone

We next isolated alignment attributable to training beyond that already present at initialization (Figure 1B). Learned features explained 43 − 80% of the noise ceiling beyond the untrained architecture, significantly above zero at every age and for every model. The contribution of learned features was similar across ages: from 4 years to adulthood it changed by less than 0.05 of the noise ceiling in every model, and none of these changes was significant (Appendix Table 1). While prior work showed that architectural priors alone can yield substantial brain-model correspondence [17; 13], our results showed that learned features add a large and age-independent share of the alignment from 4 years to adulthood.

**Table 1:** Change in training-specific alignment across age groups (Figure 1B). For each model, a straight line was fitted to the normalized values of the five age groups (4 years to adults, numbered 1–5); the slope is the average change, as a fraction of the noise ceiling, from one age group to the next. Confidence intervals and two-sided *p*-values from 10,000 bootstrap resamples of participants within each age group; Holm correction across models.

| Model (depth) | Slope per age group | 95% CI | *p* | *p*<sub>Holm</sub> |
|---|---|---|---|---|
| ResNet-50 Supervised (6) | +0.011 | [−0.032, +0.059] | .62 | 1.00 |
| ResNet-50 DINO (6) | −0.005 | [−0.052, +0.045] | .84 | 1.00 |
| ViT-B/16 Supervised (4) | +0.006 | [−0.030, +0.046] | .74 | 1.00 |
| ViT-B/16 DINO (6) | −0.009 | [−0.047, +0.033] | .66 | 1.00 |

### Model training does not show a systematic progression along the measured developmental trajectory

Prior work showed that model training progressively captures later stages of mature visual processing [18]. However, by itself this pattern does not establish that optimization follows the trajectory of human visual development. If model learning recapitulated development, successive training stages would align with progressively older neural representations. Instead, alignment with every age group emerged very early during training and remained stable thereafter (Figure 1C, left). The diagonal score was close to zero for every model (within ± 0.005 of the noise ceiling; uncorrected *p* = .21–.62, Holm *p* ≥ .83; Figure 1C, right), and its sign was not consistent across depths or seeds (Appendix Figure 5). Thus, unlike the progression through the temporal hierarchy of mature visual processing reported during model training [18], we find no evidence that optimization systematically traverses child- and adult-like representational geometries in order. This reveals a dissociation between human development and the training trajectories of the investigated models.

**Figure 5:**
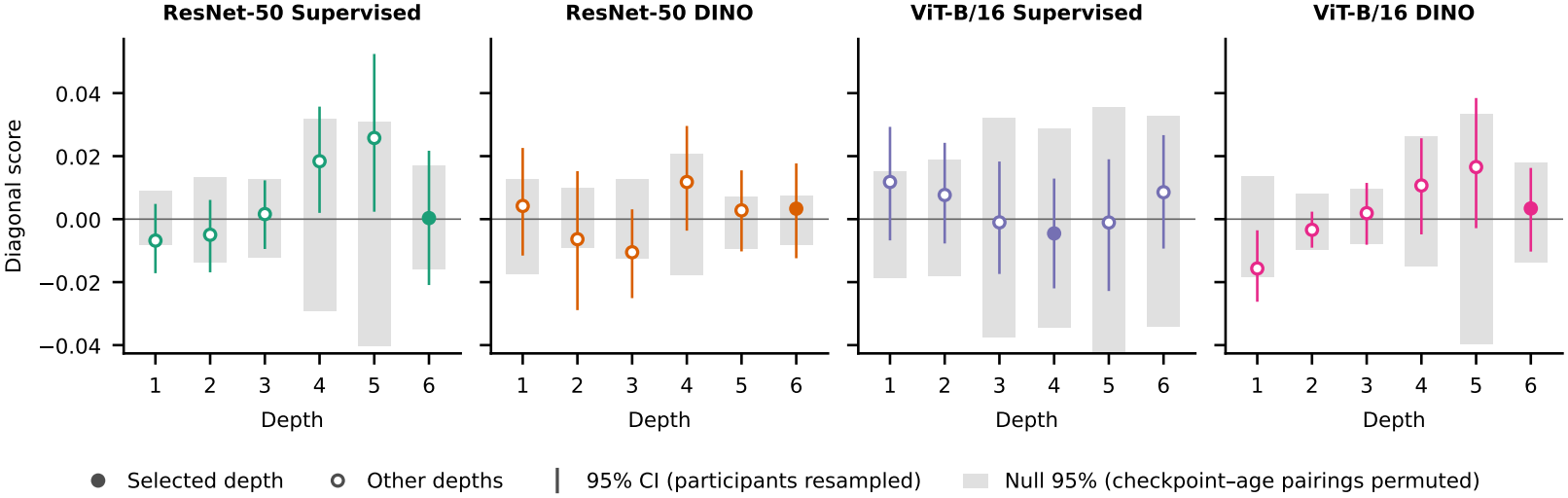
Diagonal score of every model at every depth (filled: depth shown in Figure 1C). Error bars: participant bootstrap 95% CI. Grey bars: 95% of the null over all permutations of the checkpoint-age pairing. No depth shows a score above its null band.

### Representational content is stable from 4 years onward once reliability is taken into account

To test whether alignment across ages reflects shared or developmentally changing visual features, we characterized the visual information underlying neural representational geometry across age groups (Figure 1D). At every age, categorical structure accounted for about half of the explainable geometry (48 − 61% of the ceiling) and spatial frequency for about a third (24 − 40% of the ceiling), whereas pixel intensity, color histogram, and Gabor energy contributed little. Category explained EEG geometry better than these three low-level predictors at all ages, and better than spatial frequency from 6 years onward (FDR *q* < .05). Regressing each low-level RDM out of the categorical RDM left categorical structure essentially unchanged (semi-partial correlations; Appendix Figure 7), so it is not explained by low-level differences between categories. Raw categorical correlations increased with age (from 0.09 at 4 years to 0.18 in adults; Appendix Figure 6), but this increase followed the reliability of the EEG: normalized categorical structure was similar across ages. Thus, from 4 years onward, model–brain alignment reflects correspondence with largely shared representational content rather than developmentally changing features.

**Figure 6:**
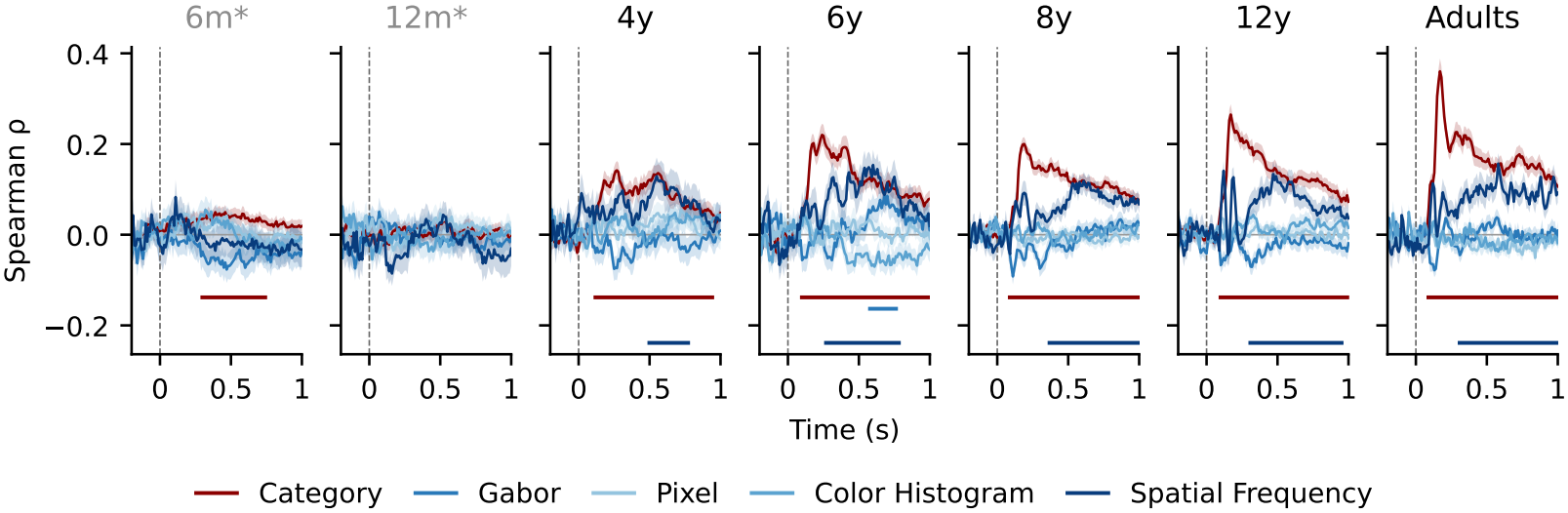
Spearman *ρ* between each participant’s EEG RDM and the categorical and low-level predictor RDMs for all seven age groups (mean ± SEM over participants). Horizontal bars mark significant clusters (one-sided sign-flip cluster test, *p* < .05), in the color of the corresponding predictor.

**Figure 7:**
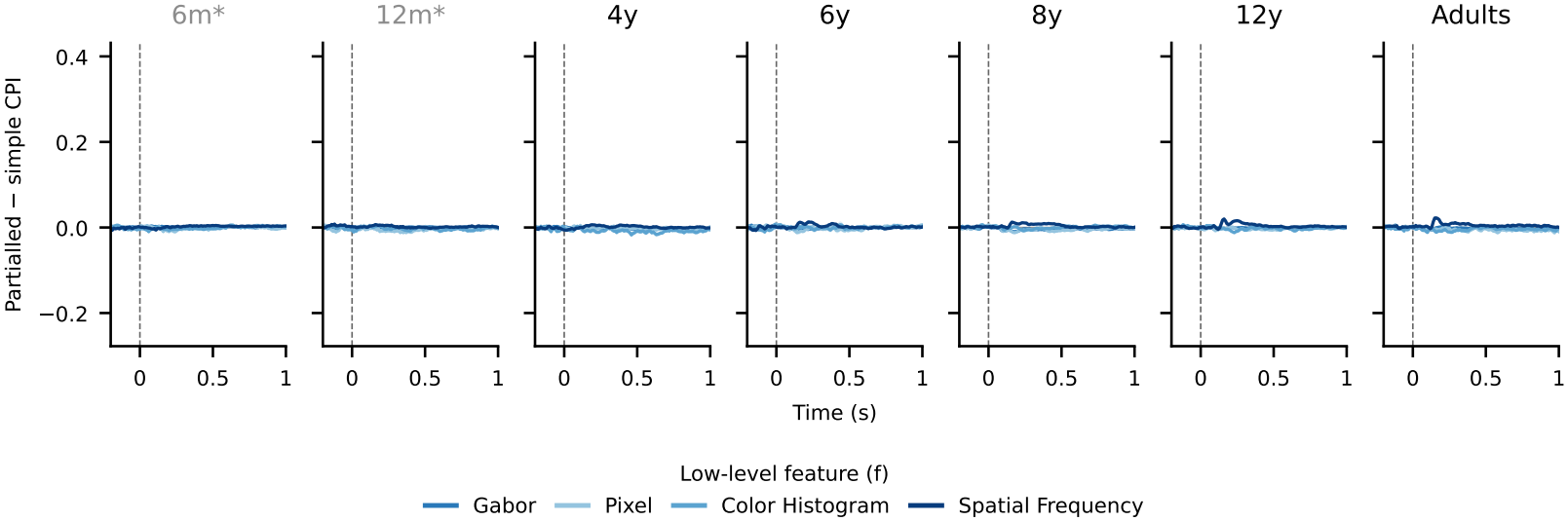
Difference between the category preference index computed from semi-partial correlations, sr(category |*f*) − sr(*f* |category), and from simple correlations, *ρ*category − *ρf* (mean ±SEM over participants), for each low-level feature *f* and all seven age groups, on the same scale as *ρ* in Fig. 6. The difference stays close to zero at every age and time point, because the categorical and low-level RDMs are nearly orthogonal (all |*r*| < 0.09).

## 4 Discussion

Sampling model states throughout training, we found no evidence that successive training stages produce child- and adult-like representational geometries in order. Instead, fully trained networks match the EEG geometry of every age group from 4 years to adulthood to a similar degree relative to EEG reliability, and this age profile was set early in training. Learned features add substantial alignment beyond architecture alone, and the categorical and low-level content of the EEG geometry was likewise similar across ages once reliability was taken into account. Several apparent developmental differences in raw measures (stronger alignment and categorical structure in adults) disappeared after normalizing by the noise ceiling, underscoring the need to account for data quality when comparing models with developing brains. Two limitations qualify these conclusions. First, the EEG geometry of 6- and 12-month-olds was not reliable in this paradigm, so the infant end of development could not be tested: this is where developmental differences in content are expected to be largest [22]. Second, the diagonal test compares only five matched ages and assumes a linear mapping from training progress to age, which limits its sensitivity. Beyond these limitations, our results raise the question of why training does not move from child-to adult-like representational geometries. One possible reason is the visual input. Standard training exposes networks to adult-quality images from the start, whereas the input to the developing visual system is shaped by visual acuity, contrast sensitivity and chromatic sensitivity that mature over time. Spatial frequency, the low-level property that most strongly structured the EEG geometry in our data, depends directly on the maturation of acuity and contrast sensitivity. Future work will test whether developmentally motivated manipulations of visual input [14] produce training trajectories that more closely resemble human visual development.

## A Brain-model alignment

Figure 2 shows the time-resolved alignment of all four fully trained models with all seven age groups. The pattern matches Figure 1A: from 4 years onward, alignment emerges around 100 ms and remains above zero throughout the epoch, with deeper layers more strongly aligned. Figure 3 compares age groups on normalized alignment, and Table 1 tests whether the training-specific alignment of Figure 1B changes with age

## B Developmental correspondence of training across model depths

Figure 4 shows the normalized age × checkpoint matrices from which the diagonal scores of Figure 1C are computed. No matrix shows the diagonal structure expected if each age group aligned best with its matched checkpoint, and the values are small (at most a few percent of the noise ceiling). Figure 5 repeats the test at every depth, showing that scores fall within their null bands and the result does not depend on the selected depth.

## C Representational content across age groups

Figure 6 shows the time courses behind Figure 1D for all seven age groups. Categorical structure is significant from about 100 ms at every age from 4 years onward, and its raw peak grows with age. Spatial frequency is the only low-level predictor that is consistently significant, later in the epoch and at a similar level across ages. Normalized by the noise ceiling, categorical structure is similar across age groups (Figure 1D), so the raw increase with age follows the reliability of the EEG. At 6 months, a weak categorical cluster reaches significance at about 0.3 − 0.75 s, but since the EEG geometry of this age group has no reliable ceiling, we do not interpret it further. Finally, we compared category with each low-level feature directly using a category preference index (CPI). The simple index, CPI*f* = *ρ*category − *ρf*, is the difference between the Spearman correlations of the EEG RDM with the categorical RDM and with the RDM of low-level feature *f* : positive values mean that category explains the EEG geometry better than *f*. Because the two predictor RDMs could share variance, we also computed a semi-partial index, sr(category |*f*) − sr(*f*| category), in which each predictor is first residualized against the other, so that only the variance unique to each is compared. The difference between the two indices stays close to zero at every age and time point (Figure 7), which is explained by the fact that the categorical and low-level RDMs are nearly orthogonal (all |*r*| < 0.09).

